# Development and validation of an open-source Hand Laterality Judgement Task for in-person and online studies

**DOI:** 10.1101/2024.10.17.618819

**Authors:** Marcos Moreno-Verdú, Siobhán M McAteer, Baptiste M Waltzing, Elise Van Caenegem, Robert M Hardwick

## Abstract

The Hand Laterality Judgement Task (HLJT) is considered a measure of the ability to manipulate motor images. The ‘biomechanical constraints’ effect (longer reaction times for hand rotations towards anatomically difficult versus biomechanically easier movements) is considered the behavioural hallmark indicating motor imagery is being used. Previous work has used diverse HLJT paradigms, and there is no standardized procedure for the task. We developed an open-source, freely available version of the HLJT in PsychoPy2, which needs no programming skills and is highly customisable. Some studies suggest responding to the HLJT with the hands may interfere with performance, which would limit practical application of the task. We examined this potential issue using in-person and online versions. For the in-person version, 40 right-footed/handed individuals performed the HLJT with their feet or bimanually (N=20 each). For the online version, 60 right-handed individuals performed the task bimanually or unimanually (N=20 each). Bayesian mixed-effect analyses quantified the evidence for and against equivalence within and between the in-person and online versions. Both versions replicated previously described behavioural phenomena, including effects of angle, hand view, and the ‘biomechanical constraints’ effect. While responding with different effectors modified overall reaction times, it did not interact with other factors analysed, and did not affect accuracy or the ‘biomechanical constraints’ effect. There was also evidence for equivalence between in-person and online bimanual groups for all measures. We conclude that this open-source, standardized HLJT protocol (available at https://osf.io/8h7ec/) can reliably detect previously identified effects and works equally well in-person or online.

## INTRODUCTION

Motor imagery is a topic of increasingly growing scientific interest. It can be defined as the mental simulation of a movement without actual physical execution (Jeannerod, 2001). Motor imagery is relevant for a variety of purposes, from fundamental neuroscientific research to applied fields in sports and rehabilitation (Ladda et al., 2021; Zhao et al., 2023). Evidence indicates that performing motor imagery is subject to inter-individual variability (Floridou et al., 2022; Guillot et al., 2008) and depends on different imagery subprocesses (Collet et al., 2011), which has led to the development of ‘motor imagery ability’ measures (Suica et al., 2022). At the behavioural level, these assessments aim to evaluate the individual’s ability to generate, maintain and manipulate motor imagery (Kraeutner et al., 2020).

The Hand Laterality Judgement Task (HLJT) has been extensively used as a measure of the ability to manipulate motor images (Parsons, 1987). In the task, individuals judge whether hand images rotated to different angles correspond to the left or right side of the body. Theoretically, most individuals will involuntarily use motor imagery to solve the task (Conson et al., 2020). That is, they will inadvertently simulate (i.e., imagine) rotating their own hand to decide the laterality. The fact that this behaviour appears unintentionally is one of the argued strengths of the HLJT over other imagery ability measures (e.g., questionnaires for imagery generation or mental chronometry for imagery maintenance), which rely on the individual’s capacity of voluntarily performing motor imagery and subjectively reporting it afterwards (Suica et al., 2022; Williams et al., 2015). The HLJT assesses so-called *implicit* motor imagery ability (McAvinue & Robertson, 2008).

Research has generally suggested that one behavioural hallmark of the HLJT may indicate whether individuals are using motor imagery to solve it: the ‘biomechanical constraints’ effect (Parsons, 1994). According to this effect, the reaction time needed to judge laterally rotated stimuli would be greater than medially rotated stimuli considering a rotation in the frontal axis, a phenomenon explained by the inherent anatomical limitations of the actual movement. While the literature is not in complete agreement with this effect solely and uniquely reflecting motor imagery-based processing (Meng et al., 2016; Vannuscorps & Caramazza, 2016), it has been widely employed as an indication of motor imagery. This is partially because the effect only appears in the HLJT as opposed to other mental rotation tasks, such as letter rotation tasks (Bek et al., 2022, p. 20; Mibu et al., 2020), which suggest the task elicits a different cognitive strategy, probably based on egocentric processing.

While the HLJT has been widely used in psychological, neuroscientific, and clinical research, it has been substantially less used in applied contexts such as rehabilitation, sports science, or online studies. Moreover, there is no standardized version of the task, and previous studies have used a wide variety of experimental paradigms (e.g. different angles of rotation, views of the hand, etc.). Furthermore, researchers have generally implemented this task in software which are typically not prepared for running both in-person and online studies. New developments in open-source stimulus presentation software can overcome both issues while permitting the experimental setup to be flexible.

One of the peculiarities of the HLJT, which might explain its limited use in applied circumstances and online studies is that if participants respond using their own hands, a possible confound with task performance might arise, in the form of an interference or priming effect (Cocksworth & Punt, 2013). This may occur because the same effector is being used to process the stimulus and provide a response, potentially modifying how information is processed while the response is being prepared. Thus, verbal (Ionta et al., 2007) or foot (Brady et al., 2011) responses have been used to avoid this potential issue. We note, however, that studies which have used the HLJT with manual responses have broadly replicated the main behavioural phenomena of this paradigm, including the presence of the ‘biomechanical constraints’ effect (Bek et al., 2022; Hudson et al., 2006; Kraeutner et al., 2019; Saimpont et al., 2009). In fact, the assertion that hand response modes modify behaviour in the HLJT has only been directly tested in one previous study (Cocksworth & Punt, 2013). Further evidence in this regard is necessary to expand the applicability of the task to non-laboratory settings, where the individual could potentially need to respond with one or two hands depending on the specific situation (e.g., in case of unilateral motor disturbances or verbal communication disorders, or online studies).

The present study therefore had two central aims. Our first aim was to develop a standardized version of the HLJT as open-source software that is freely available. We publicly share both local and online running versions of the paradigm we used for this study (https://osf.io/8h7ec/). The task allows a high level of customizability and does not require extensive programming skills to be set up, with the ultimate goal of paving the way for its use in non-laboratory contexts. We ‘validated’ the in-person and online versions against each other using Bayesian analyses and equivalence tests, and investigated whether both versions could replicate the well-established behavioural phenomena in this task. The second goal was to test the hypothesis that manual responses interfere with performance in the HLJT, as this would facilitate future use in applied or online contexts. In the in-person version we compared a bimanual response mode against a foot response mode in terms of accuracy, reaction time and, critically, the ‘biomechanical constraints’ effect. We predicted that manual responses could modify task performance in terms of accuracy or reaction time, but not the biomechanical effect. Importantly, even in the presence of small discrepancies between response modes, we posited that meaningful differences would not emerge, and that responding with the hands would be practically equivalent than responding with the feet. In the online version, we compared the equivalence between bimanual and unimanual response modes, as the latter may be necessary for clinical applications.

## METHODS

### Study design and participants

This study used a mixed design, comparing measurements within- and between-subjects and in-person or online versions of the HLJT. All study procedures followed the Declaration of Helsinki (2013 revision) and were approved by the Ethics Committee of the Institute for Research in the Psychological Sciences, UCLouvain (N°: 2024-21). All participants were financially compensated for their time (€10/h).

Overall, 100 right-handed healthy individuals aged 18-35 years, with normal or corrected-to-normal vision and no history of neurological damage participated. Participants were identified as right-handed by self-assessment. All participants were assessed with the Movement Imagery Questionnaire-3 (MIQ-3) to determine the ability to generate motor imagery as total, kinesthetic, internal visual and external visual imagery scores (Williams et al., 2012).

#### Setting and participants in-person version

Forty right-handed individuals participated (see Table 1 for demographics). All participants were also self-identified as right-footed by answering the question “If you were to kick a ball would you do it with your right leg?”, as previous work has established this as an appropriate way to assess leg dominance (van Melick et al., 2017). Participants were recruited via public advertisements, and they were assessed at UCLouvain (Belgium). Participants were randomly allocated to 2 different groups which only differed in the Response Mode. A computer random sequence generator with 1:1 assignment was used. The “Foot” group (N = 20) responded to the HLJT with their feet, whereas the “Bimanual” group (N = 20) responded with their hands (see Response Modes for details).

**Table 1.** Participants’ characteristics for the in-person and online Hand Laterality Judgement Task versions according to their ps, including the Bayes Factors in favour of the null hypothesis of no differences between the groups (BF_01_).

| Characteristic | In-person version |  |  | Online version |  |  |  |
| --- | --- | --- | --- | --- | --- | --- | --- |
|  | Foot<br>(N = 20) | Bimanual<br>(N = 20) | Bayes Factor for<br>null (BF<sub>01</sub>) | Left-Hand<br>(N = 20) | Bimanual<br>(N = 20) | Right-Hand<br>(N = 20) | Bayes Factor for<br>null (BF<sub>01</sub>) |
| **Age, mean ± SD** | 24.80 ± 3.65 | 24.25 ± 3.21 | 2.92 | 26.20 ± 4.97 | 25.70 ± 3.63 | 26.05 ± 3.82 <sup>1</sup> | 6.94 |
| **MIQ-3, mean ± SD** |  |  |  |  |  |  |  |
| Total score | 68.40 ± 8.91 | 66.95 ± 7.44 | 2.86 | 61.30 ± 9.89 | 63.05 ± 11.75 | 61.79 ± 12.17 | 6.62 |
| Kinesthetic | 21.85 ± 3.67 | 22.05 ± 4.15 | 3.21 | 20.20 ± 4.26 | 20.90 ± 4.33 | 21.10 ± 3.93 | 6.15 |
| Internal Visual | 22.65 ± 3.80 | 22.30 ± 3.99 | 3.14 | 20.65 ± 3.88 | 21.05 ± 4.22 | 19.84 ± 4.68 <sup>2</sup> | 5.46 |
| External Visual | 23.90 ± 2.94 | 22.60 ± 3.55 | 1.73 | 20.45 ± 3.98 | 21.26 ± 3.98 <sup>3</sup> | 20.95 ± 4.25 <sup>3</sup> | 6.26 |
| **Sex, n (%)** |  |  | 1.18 |  |  |  | 6.29 |
| Female | 12 (60%) | 16 (80%) | - | 9 (45%) | 11 (55%) | 9 (45%) | - |
| Male | 8 (40%) | 4 (20%) | - | 11 (55%) | 9 (45%) | 11 (55%) | - |

<sup>1</sup> Data showed from N=19 participants. <sup>2</sup> Data showed from N=19 participants. <sup>3</sup> Data showed from N=19 participants.
Note: BF<sub>s</sub> are interpreted following established benchmarks, considering 'no' evidence, 'anecdotal', 'moderate', 'strong', 'very strong' and 'extreme' evidence if the BF was ≈1, 1-3, 3-10, 10-30, 30-100 and >100, respectively (Andraszewicz et al., 2015; Jeffreys, 1998). Bayes factors are presented in relation to the null (BF<sub>01</sub>) hypothesis.

#### Setting and participants online version

Sixty right-handed individuals participated (see Table 1 for demographics). In order to verify the task could be completed online, the experiment was conducted fully remotely, and participants were recruited via the crowdsourcing platform Prolific (https://www.prolific.com/). Participants were randomly allocated to 3 different groups (N = 20 each) that completed the task in either a “Left Hand”, “Bimanual” or “Right Hand” condition (see Response Modes for details).

### General procedure, task, and stimuli

#### General procedure

Experimental procedure is shown in Fig. 1A and Fig. 1B and was common for both versions. First, participants were assessed using an electronic version of the MIQ-3. The cross-culturally adapted English, Spanish or French versions were used depending on the native language of the participant (Robin et al., 2020; Trapero-Asenjo et al., 2021). Participants completed the questionnaire on their own. In the in-person version, the experimenter clarified any doubts or corrected the participants only if they asked or if they did not perform the movements as intended (e.g., if they performed a completely different movement or did the movement with the contralateral limb). In the online version, no clarifications were made.

**Figure 1.**
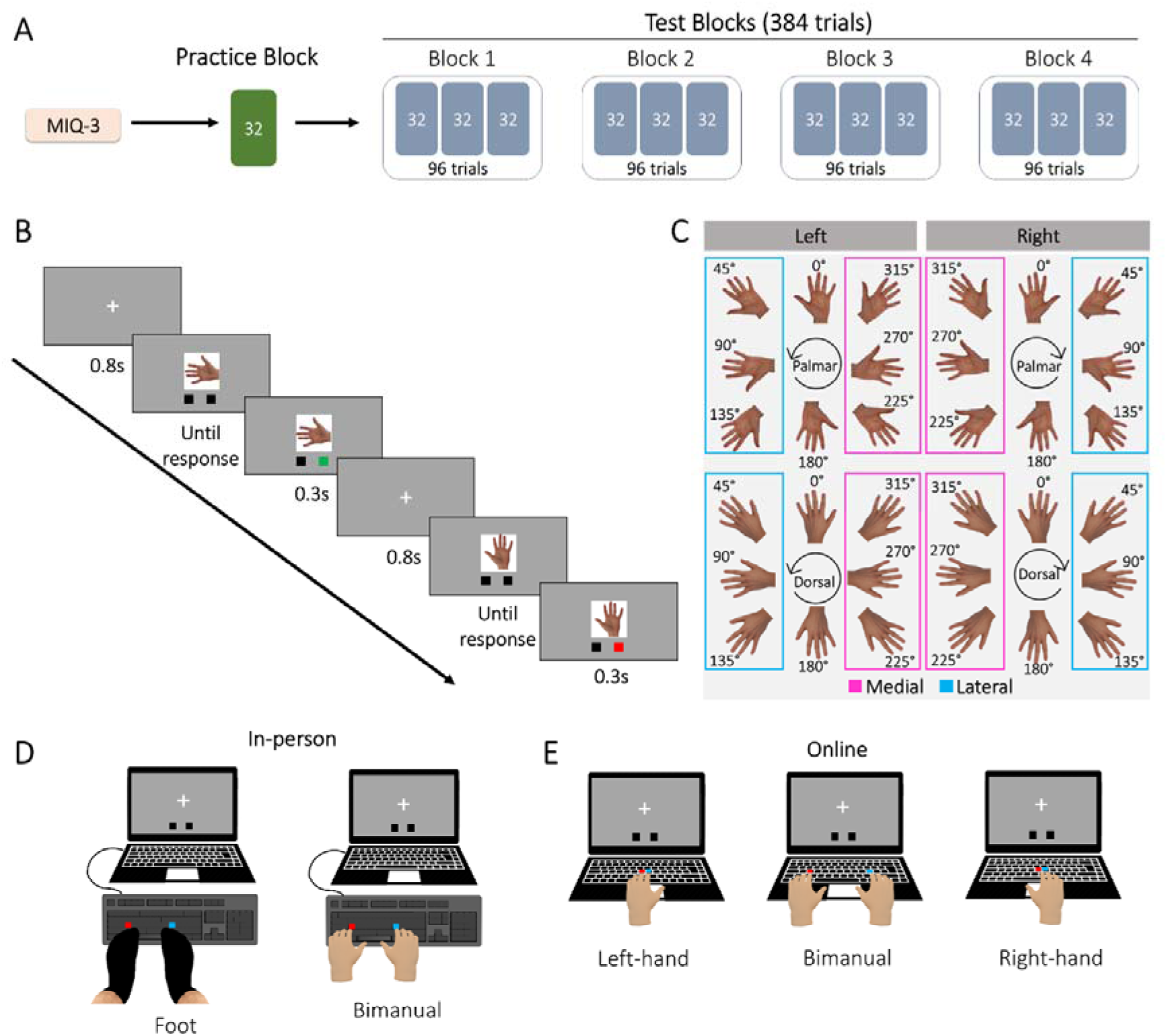
Experiment overview. **Panel A** shows the overall structure of the experiment. Participants started by completing the electronic version of the Movement Imagery Questionnaire-3 (MIQ-3) before doing the Hand Laterality Judgement Task (HLJT), which consisted in 1 practice block and 4 test blocks. **Panel B** shows the experimental stimuli used Laterality (right/left) and rotational angle (0° to 315° in increments of 45°) were used to code medial and lateral directions according to the anatomical position, which were later used for the ‘biomechanical constraints’ effect analysis. **Panel C** shows an example of trials from the HLJT, with visual feedback (green for correct responses, red for incorrect responses). **Panels D and E** show all the possible Response Modes used in in-person (panel D) and online (panel E) versions of the HLJT. The Foot and Bimanual groups used the keys ‘S’ and ‘L’ of the keyboard (left and right respectively). The Foot group responded on an external keyboard placed on the floor from which all keys but ‘S’ and ‘L’ had been removed. The Bimanual group used the index finger of their corresponding hand on the same external keyboard placed on the desk. The Bimanual group in the online version used the same paradigm as in-person, but on the computer’s keyboard. The Left-hand and Right-hand groups used the index and middle fingers placed over the ‘G’ and ‘H’ keys (left and right respectively) on the computer’s keyboard.

Participants were assessed by the HLJT, whereby they judged whether images of hands that had been rotated depicted a right or left hand. Two experiments were created and run in PsychoPy2 version 2023.2.3 (Peirce et al., 2019). A ‘local’ experiment was used for the in-person version, whereas for the online version this was output to a PsychoJS experiment to be run online via PsychoPy2’s associated webpage Pavlovia (https://pavlovia.org/) (Bridges et al., 2020). The files to run both experiments are freely accessible at https://osf.io/8h7ec/.

All participants were instructed to respond as quickly and accurately as possible through standardized on-screen instructions. The instructions were available in English, French or Spanish and had been reviewed by two native speakers in each case to ensure equivalence. In the in-person version, participants sat at a comfortable distance to the experiment computer (a 14-inch Dell laptop, 16gb RAM, 13^th^ Gen i5-1345U, 1.6GHz, refresh rate = 60Hz) in a quiet room. In the online version, participants completed the experiment on their own computer, and were instructed to complete the task in a quiet room without distractions.

#### Task and experimental stimuli

The task and experimental stimuli were identical in both versions. The experimental stimuli were real hand pictures as shown in Fig. 1C. Images of left hands were right-hand images that had been mirror reversed. Left or right images were presented in 8 rotational angles in the frontal axis (0°, 45°, 90°, 135°, 180°, 225°, 270°, 315° for right hands in the clockwise direction, and the opposite for left hands), and 2 possible views (palmar or dorsal). In total, 32 unique stimuli were used.

Each stimulus was preceded by a small fixation cross in the centre of the screen for 800ms. Stimuli were presented in the centre (PsychoPy dimensions: 0.45 x 0.45 units) and remained until a response was made. Visual feedback following each stimulus was provided for 300ms via two small boxes (PsychoPy dimensions: 0.07 x 0.07 units) placed at the bottom of the screen; the corresponding box turned green if a response was correct, and red if incorrect (Fig. 1B). Participants were allowed to familiarise themselves with the task and stimuli in a practice block with 1 repetition per unique stimulus (i.e., 32 trials) followed by 4 test blocks with 96 trials each (3 repetitions per unique stimulus). In each test block, stimuli were randomly presented in sub-blocks of 32 trials with 1 repetition per unique stimulus in each sub-block, to minimize the likelihood of the same stimulus appearing more than twice in a row. Only test blocks were used for the analyses, resulting in a total of N = 384 trials per participant. Breaks between blocks were allowed, with a minimum duration of 10 seconds and no maximum limit.

### Response Modes across versions

Participants in both versions of the HLJT responded with one of the Response Modes depicted in Fig. 1D-E. All participants were given standardized, on-screen instructions which differed only according to the corresponding Response Mode.

In the in-person version (Fig. 1D), participant responses were recorded using an external QWERTY USB-keyboard (dell). All buttons were removed from this keyboard apart from the ‘S’ and ‘L’ keys, which were used to provide responses to left- and right-hand stimuli, respectively. This procedure allowed us to record both hand and foot responses with the same device. For the “Foot” Response Mode the keyboard was placed on the floor inside a custom-built wooden frame to help maintain its position. Participants sat at a desk and placed their left foot in contact with the ‘S’ key, and their right foot in contact with the ‘L’ key. They were instructed to maintain the position of the feet as stable as possible, with the heels in contact with the wooden frame, and did not wear shoes. Participants were allowed to adjust the positioning of their legs to optimise comfort, although their knees were always flexed at approximately 90°. They were asked to place their hands over their thighs and under the desk (i.e., without seeing them), and asked not to move them while performing the task. In the “Bimanual” mode, the keyboard was placed on the surface of the desk, and participants sat with their left index finger in contact with the ‘S’ key, and their right index finger in contact with the ‘L’ key. As such, participants could see their hands during the task.

In the online version (Fig. 1E), participants were instructed to complete the task either unimanually (with only the left or right hand), or bimanually. Participants in the “Bimanual” mode responded as in the in-person version, but on their own computer’s keyboard. In the “Left-hand” and “Right-hand” Response Modes (only online), participants were instructed to respond with the index and middle fingers of the respective hand on their computer’s keyboard. The keys in these two modes were the same and were centred on the keyboard (‘G’ for left images and ‘H’ for right images) to maintain relative spatial congruence in the required responses(Waltzing et al., 2024).

### Statistical analysis

All analyses were performed in R version 4.3.3 (R Core Team 2024). Scripts and data are freely available at https://osf.io/8h7ec/. As a large part of the predictions for this study were focused on considering the evidence in favour of the null hypotheses or practically equivalent differences, a Bayesian framework was used. This allowed us to obtain evidence for and against the null/alternative hypothesis. Therefore, all primary results are reported from the posterior distribution as means and 95% Credible Intervals (95%CrI). Additionally, descriptive statistics from sample data (mean and SD) are presented in Supplementary Materials.

Accuracy and reaction time were analysed separately. First, trials where reaction time was <300MS or >3,000ms, likely reflecting anticipatory responses and no engagement with the task, respectively, were discarded. For accuracy, the proportion of correct responses for each unique stimulus for each participant, after averaging across repetitions, was obtained. For reaction time, the mean time for each unique stimulus was considered only for the trials with correct responses. The measure of the ‘biomechanical constraints’ effect was the difference between the mean reaction time of medially vs. laterally rotated stimuli (Fig. 1C).

Separate models for the in-person and online datasets were first created and analysed. Then, data from the two Bimanual groups (i.e., in-person and online) were combined and analysed, to further validate the equivalence of online version of the task with its in-person counterpart.

Statistical modelling was performed with multilevel models (i.e., mixed-effects models), considering by-participant random intercepts using the ‘brms’ package (Bürkner, 2017). These models were applied with a Gaussian identity likelihood for reaction time and a zero-and-one inflated Beta likelihood for accuracy (as it allowed the proportion of correct responses to be in the range 0-1 *including* both 0s and 1s in the distribution) (Liu & Eugenio, 2018). The full model was always of type *‘y ∼ Group*Angle*Laterality*View + (1|Participant)’*, where Group was a between-subject factor representing different Response Modes (for comparisons within the same version of the study) or setting (for comparisons between the in-person and online versions).

For the ‘biomechanical constraints’ effect, a subset of the data considering only trials where this effect could be present (i.e., rotations in the lateral (45°, 90°, 135°) and medial (225°, 270°, 315°) directions) was used. The full model was *‘Reaction Time ∼ Group*Direction*Laterality*View + (1|Participant)’*, where the effect of interest was the Group * Direction interaction, or further higher-order interactions including these factors. The biomechanical constraints effect was therefore analysed by collapsing across the three rotation angles for each direction.

All models were run with weakly informative priors for all the beta parameters. For reaction time, this was a normal distribution of mean = 0 and SD = 300ms, whereas accuracy models were run with a normal distribution of mean = 0 and SD = 1 on the logit scale. The same applied for the ‘biomechanical constraints’ models for reaction time. Default (uniform) priors were used for the remaining parameters. All models were run with 10 chains of 5,000 iterations each (1,000 warmup iterations per chain) for an overall post-warmup of 40,000 iterations to inform the posterior distribution. Model fit was assessed by visually inspecting posterior predictive checks and trace plots, and R^2^ values at convergence (R^2^ < 1.01 was considered appropriate). More details on the Bayesian models used can be found in Supplementary Materials.

For hypothesis testing, we used Bayesian Model Averaging to obtain the inclusion Bayes Factor (BF_inc_) for including each given effect or interaction, or against (BF_01_) including it (Hinne et al., 2020). BF_inc_ were obtained by bridge sampling (Gronau et al., 2017) using the ‘bayestestR’ package (Makowski et al., 2019). If evidence was found for a given effect of Response Mode, or an interaction of Response Mode and other factors, post-hoc comparisons were performed as equivalence tests considering a Region of Practical Equivalence (ROPE) of 0.1*SD of the outcome variable (Kruschke, 2018). Equivalence tests were run to examine evidence in favour of the null hypothesis of equivalence (BF_01_), or the alternative hypothesis of non-equivalence (BF_10_), always with equal-prior factor coding (Morey & Rouder, 2011). For post-hoc comparisons not involving Response Mode, point-null hypothesis testing was used. BFs for post-hoc comparisons were obtained via the Savage-Dickey density ratio. All BFs were interpreted following established benchmarks, considering ‘no’ evidence, ‘anecdotal’, ‘moderate’, ‘strong’, ‘very strong’ and ‘extreme’ evidence if the BF was ≈1, 1-3, 3-10, 10-30, 30-100 and >100, respectively (Andraszewicz et al., 2015; Jeffreys, 1998). Bayes factors are presented in relation to the null (BF_01_) or alternative (BF_10_) hypothesis.

## RESULTS

### In-person version

Overall, 1.35 ± 0.27% (mean ± SD) of trials in the HLJT were rejected due to extremely short (<300ms) or long (>3000ms) reaction time. The mean time to complete the task was 17m 3s ± 1m 26.4s. There was evidence against including a main effect of Response Mode or interactions between this factor and other terms for reaction time, accuracy and the ‘biomechanical constraints’ effect. Therefore, we first present results collapsing across groups, but we also report them separately (see below for details).

#### Reaction time

Overall, 13,109 valid trials were analysed. There was extreme evidence for including the main effects of Angle, and View, and an interaction between them (BF_inc_ = 3.21×10^138^, BF_inc_ = 3.17×10^17^ and BF_inc_ = 5.8×10^17^, respectively). Evidence against including a main effect of Laterality, or two-way and three-way interactions was very strong (BF_01_ > 35.71). There was evidence against including interactions of these factors with Response Mode.

The main effect of Angle was explained by an increase in reaction time with stimulus rotation (Fig. 2A, see Table 2 for pairwise comparisons), reaching the longest reaction time at the maximum absolute rotation (180°). The main effect of View was explained by palmar views (mean = 1038.69ms [979.97, 1096.95] 95%CrI) being faster than dorsal views (mean = 1066.09ms [1007.54, 1124.87]), with inconclusive evidence for the difference not being 0 (difference = -27.4ms [-57.8, 6.71], BF_10_ = 1.16).

**Figure 2.**
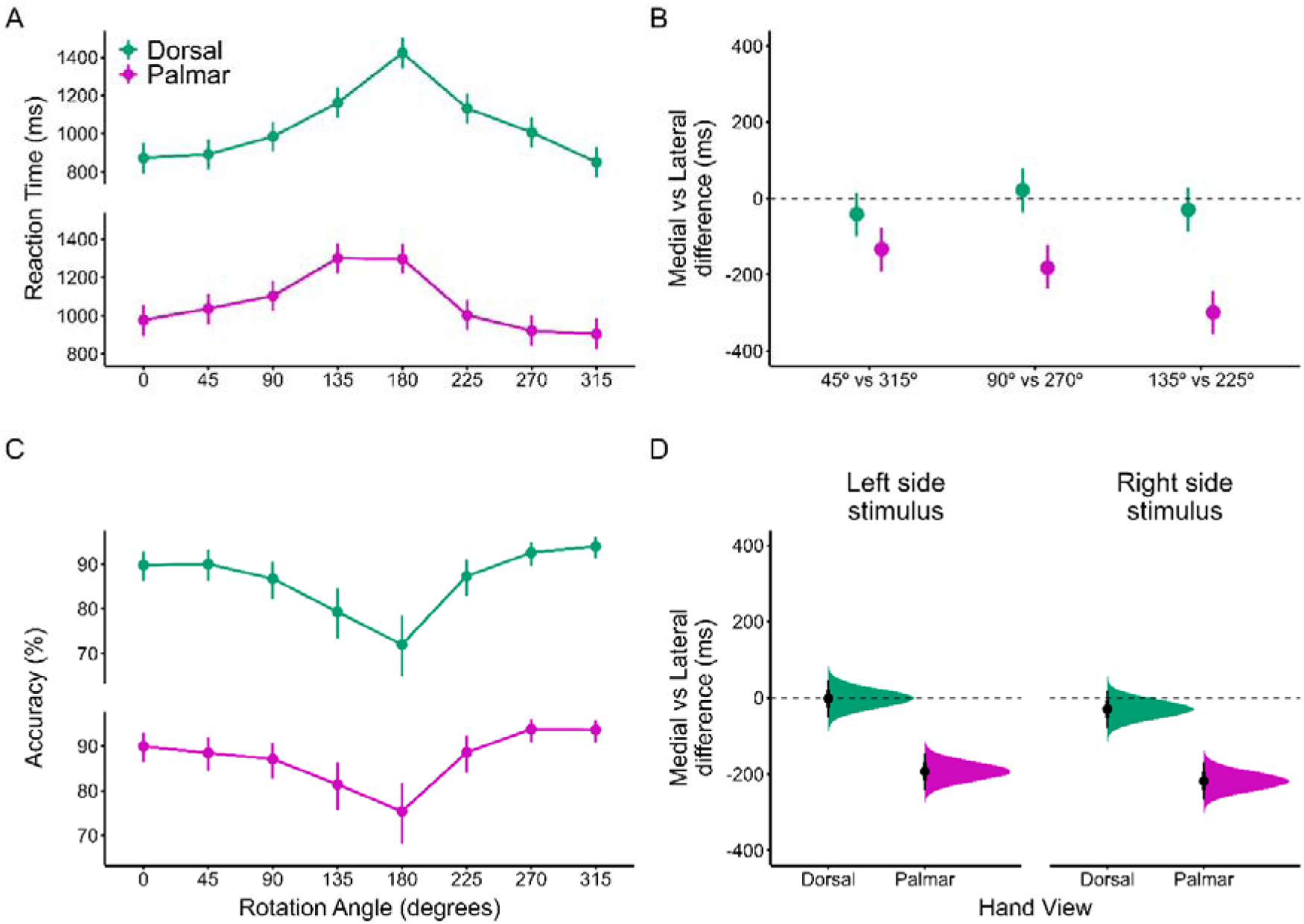
Results of the in-person (N=40) version of the Hand Laterality Judgement Task. **Panels A and B** show the Reaction Time and Accuracy measures by Rotation Angle and View. **Panel C** shows the ‘biomechanical constraints’ effect (medial vs lateral difference in milliseconds) for the corresponding pairs of angles. **Panel D** shows the ‘biomechanical constraints’ effect collapsed across angles, split by View and Laterality.

**Table 2.** Bayes Factors in favour of the null hypothesis of a zero difference (BF_01_; in bold) or the alternative (BF_10_) hypothesis for the pairwise comparisons between neighbouring Angles for Accuracy (%) and Reaction Time (milliseconds) in the Hand Laterality Judgement Task.

| Comparison | In-person version |  | Online version |  |
| --- | --- | --- | --- | --- |
|  | Reaction Time | Accuracy | Reaction Time | Accuracy |
| 0° vs 45° | **$BF_{01} = 2.48$** | **$BF_{01} = 9.35$** | **$BF_{01} = 1.35$** | **$BF_{01} = 4.74$** |
| 45° vs 90° | $BF_{10} = 59.11$ | **$BF_{01} = 1.73$** | $BF_{10} = 3.13 \times 10^6$ | **$BF_{01} = 2.26$** |
| 90° vs 135° | $BF_{10} = 1.49 \times 10^9$ | $BF_{10} = 1.39$ | $BF_{10} = 1.44 \times 10^6$ | $BF_{10} = 111.46$ |
| 135° vs 180° | $BF_{10} = 2.45 \times 10^4$ | $BF_{10} = 21.60$ | $BF_{10} = 5.55 \times 10^4$ | **$BF_{01} = 8.06$** |
| 180° vs 225° | $BF_{10} = 2.39 \times 10^{18}$ | $BF_{10} = 5.96 \times 10^4$ | $BF_{10} = 7.2 \times 10^{15}$ | $BF_{10} = 1.96 \times 10^5$ |
| 225° vs 270° | $BF_{10} = 6.91 \times 10^3$ | $BF_{10} = 227.97$ | $BF_{10} = 2.01 \times 10^5$ | **$BF_{01} = 2.63$** |
| 270° vs 315° | $BF_{10} = 265.02$ | **$BF_{01} = 9.8$** | $BF_{10} = 879.18$ | **$BF_{01} = 6.8$** |

The interaction between Angle and View suggested the presence of the ‘biomechanical constraints’ effect (Fig. 2B). Post hoc-analyses examined this separately for palmar and dorsal views. Data for palmar views was consistent with a ‘biomechanical constraints’ effect, with the lateral rotations (45-135°) being slower than their medial counterparts (225-315°). Pairwise comparisons showed extreme evidence for a difference in each comparison (45° vs 315°: BF_10_ = 840.01; 90° vs 270°: BF_10_ = 1.48×10^5^; 135° vs 225°: BF_10_ = 1.91×10^11^). By contrast, there was no evidence for a ‘biomechanical constraints’ effect for dorsal views, with moderate evidence in favour of the null hypothesis for each comparison (45° vs 315°: BF_01_ = 4.17; 90° vs 270°: BF_01_ = 8.70; 135° vs 225°: BF_01_ = 7.04).

#### Accuracy

Overall, 15,153 valid trials were analysed. Only extreme evidence for including a main effect of Angle was found (BF_inc_ = 1.47×10^24^; Fig. 2C), explained by a decrease in accuracy with stimulus rotation, reaching its minimum value at the highest absolute rotation (180°). Pairwise comparisons are shown in Table 2. Extreme evidence against including all the remaining main effects and interactions was found (BF_01_ > 6849.32).

#### Biomechanical constraints effect

When considering only trials that could be affected by biomechanical constraints (i.e. pooling data for ‘lateral’ and ‘medial’ stimuli while excluding 0° and 180° rotations), a total of 10,053 valid trials were available for analysis. Extreme evidence for including a main effect of Direction (BF_inc_ = 5.1×10^23^), along with a Direction x View interaction (BF_inc_ = 7.49×10^10^) were found. Moderate evidence for including a main effect of Laterality (BF_inc_ = 6.39) and strong evidence for including a Direction x Laterality interaction was found (BF_inc_ = 11.72). Strong evidence for including a three-way Direction x View x Laterality interaction (BF_inc_ = 15.38) was also found.

The main effect of Direction was explained by lateral directions being slower than medial directions, consistent with the ‘biomechanical constraints’ effect (lateral: 1079.38ms [1019.64, 1137.92]; medial: 968.66ms [910.30, 1027.83]; difference = - 111ms [-136, -86.8], BF_10_ = 3.55×10^6^). This effect was critically conditioned on View (Fig. 2D), as for dorsal views, there was strong evidence in favour of the null hypothesis of the difference being 0 (difference = -15.4ms [-50.7, 19.1], BF_01_ = 12.99), whereas for palmar views there was extreme evidence for a difference not being 0 (difference = -205.8ms [-240.6, -172.3], BF_10_ = 1.55×10^10^). Stimulus laterality weakly conditioned the effect, as the difference was greater for right hands than for left hands (right = -124ms [-158, -88.7], BF_10_ = 4.62×10^6^; left = -97.4ms [-132, -62.7], BF_10_ = 6.12×10^3^). The three-way interaction revealed more marked ‘biomechanical constraints’ for right palmar (difference = -218.56ms [-267.8, -170.3], BF_10_ = 1.1×10^8^) than left palmar (difference = -193.08ms [-243.2, 147.0], BF_10_ = 1.56×10^7^), whereas for dorsal views, neither side showed evidence for a biomechanical effect (right = - 29.54ms [-77.0, 19.9], BF_01_ = 7.52; left = -1.57ms [-50.4, 47.8], BF_01_ = 14.93).

#### Effects of Response Mode

The main results are shown in Fig. 3. For reaction time, there was very strong evidence against including a main effect of Response Mode (BF_01_ = 32.26), and very strong to extreme evidence against including two-way, three-way, or four-way interactions (BF_01_ > 83.33, BF_01_ > 1.46×10^5^ and BF_01_ > 1.32×10^14^, respectively). The Foot Group (mean = 979.47ms [898.86, 1061.71]) was faster than the Bimanual Group (mean = 1125.32ms [1044.06, 1206.25]), with moderate evidence in favour of non-equivalence (BF_10_ = 3.78; ROPE = 0 ± 31.27ms).

**Figure 3.**
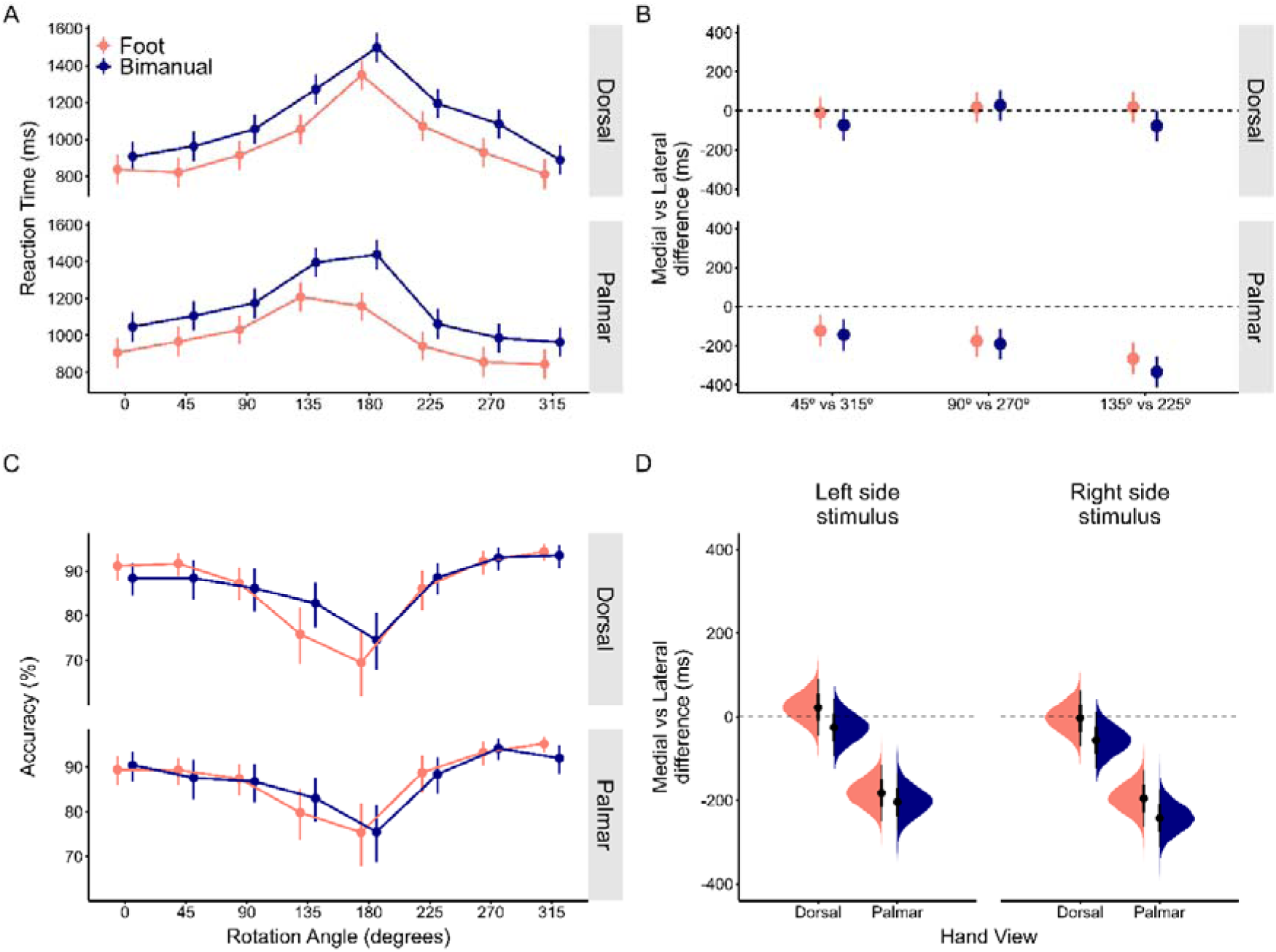
Comparison of the Foot and Bimanual Response Modes (N=20 each) of the in-person version of the Hand Laterality Judgement Task. **Panels A and B** show the Reaction Time and Accuracy measures by Rotation Angle and View. **Panel C** shows the ‘biomechanical constraints effect’ (medial vs lateral difference in milliseconds) for the corresponding pairs of angles. **Panel D** shows the ‘biomechanical constraints’ effect collapsing across angles but splitting by View and Laterality.

For accuracy, extreme evidence against including a main effect of Response Mode was found (BF_01_ = 5.68×10^8^), as well as interactions including this factor (BF_01_ = 3.64×10^8^). Accuracy was comparable between the Foot Group (mean = 86.66% [82.39, 90.33]) and the Bimanual Group (mean = 87.06% [82.6, 90.88]), with strong evidence in favour of equivalence (BF_01_ = 20.83; ROPE = 0 ± 3.42%).

For the ‘biomechanical constraints’ effect model, evidence for including a Response Mode x Direction interaction was inconclusive but favoured the null hypothesis (BF_01_ = 1.46). The ‘biomechanical constraints’ effect was present in both groups (Foot = - 89.5ms [-124, -55.5], BF_10_ = 3.58×10^3^; Bimanual = -132ms [-166, -96.9], BF_10_ = 6.83×10^5^), with moderate evidence for equivalence (BF_01_ = 4.9). Evidence against including interactions with View or Laterality was moderate (BF_01_ = 5.29 and BF_01_ = 4.52, respectively). Evidence against three-way interactions involving Response Mode, or a four-way interaction, was strong to extreme (BF_01_ > 52.63 and BF_01_ = 30581.04, respectively).

### Online version

Overall, 4.36 ± 3.04% of trials were rejected due to extremely short (<300ms) or long (>3000ms) reaction time. The mean time to complete the task was 22m 22s ± 5m 40s. Evidence against including an effect of Response Mode was found in terms of accuracy and the ‘biomechanical constraints’ effect, as well as evidence against higher-order interactions in these models. For reaction time, evidence for including a main effect of Response Mode, and a Response Mode by View interaction was found (see below for details).

#### Reaction time

Overall, 19,544 valid trials were analysed. Extreme evidence for including the main effects of Angle (BF_inc_ = 2.76×10^187^), View (BF_inc_ = 9×10^32^) and Laterality (BF_inc_ = 551.26), for their two-way interactions (Angle x View: BF_inc_ = 1.69×10^33^, Angle x Laterality: BF_inc_ = 1030, Laterality x View: BF_inc_ = 1030) and for a three-way interaction (BF_inc_ = 1380) was found.

The effect of Angle was explained by an increase in reaction time with stimulus rotation (Fig. 4A, see Table 2 for pairwise comparisons), reaching the longest reaction time at the maximum absolute rotation (180°). The effect of View was explained by dorsal stimuli showing longer reaction time than palmar stimuli (dorsal: 1223.21ms [1148.21, 1298.78]; palmar: 1178.04ms [1101.81, 1253.97]; BF_10_ = 3200). The effect of Laterality was explained by left stimuli showing longer reaction time compared to right stimuli (left: 1229.23ms [1154.24, 1306.47]; right: 1172.02ms [963.0, 1248.16]; BF_10_ = 5.49×10^5^). Critically, the interaction between Angle and View was consistent with the ‘biomechanical constraints’ effect (Fig. 4B). Post-hoc analysis for palmar views indicated that lateral rotations (45-135°) were slower than their medial counterparts (225-315°). Pairwise comparisons showed extreme evidence for the difference not being 0 at each rotation angle (45° vs 315°: BF_10_ = 263.55; 90° vs 270°: BF_10_ = 9.67×10^8^; 135° vs 225°: BF_10_ = 3.46×10^14^). Further post-hoc analysis identified that this effect did not appear for dorsal views, with anecdotal to strong evidence in favour of the null hypothesis (45° vs 315°: BF_01_ = 2.58; 90° vs 270°: BF_01_ = 10; 135° vs 225°: BF_01_ = 13.51).

**Figure 4.**
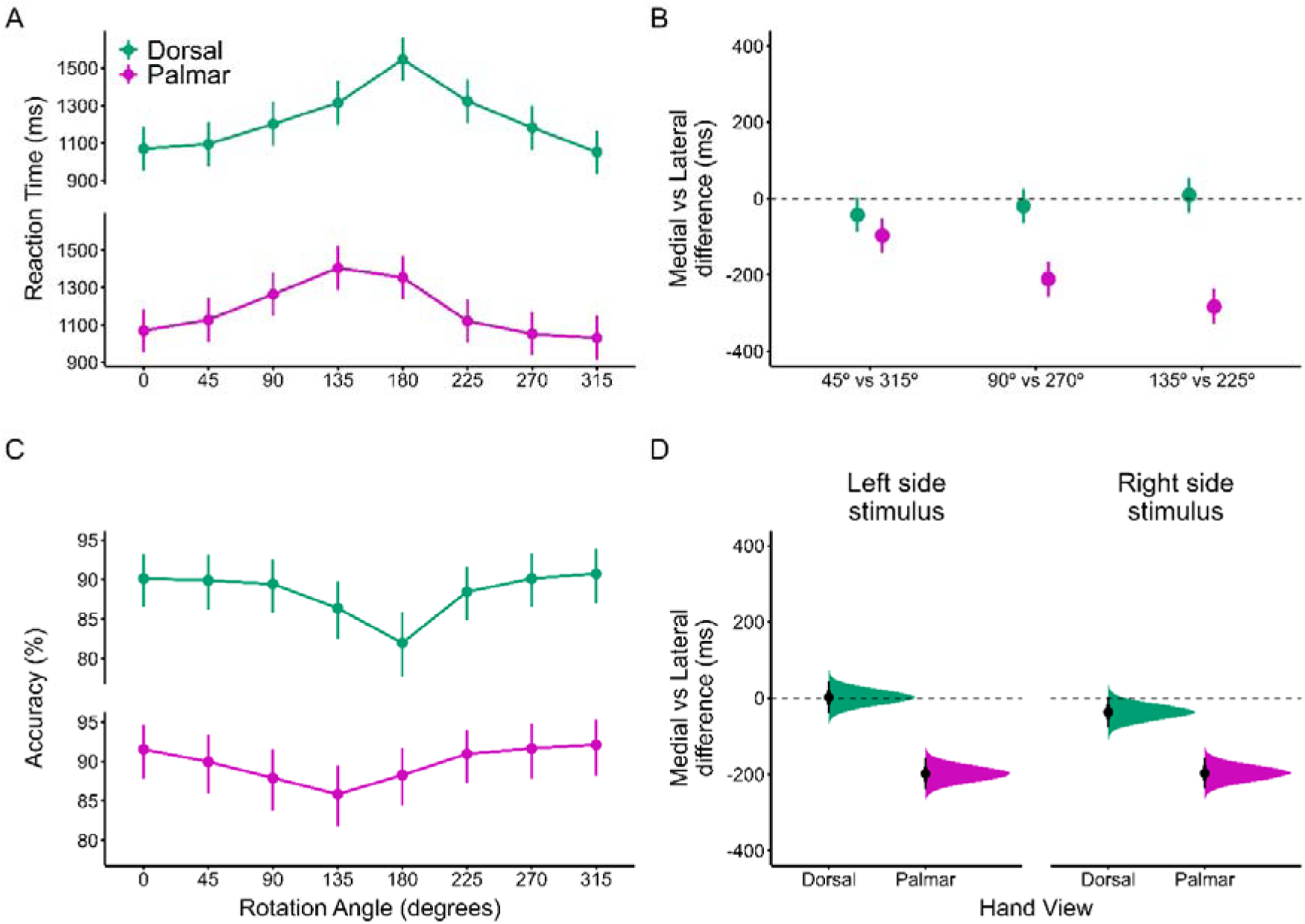
Results of the online (N=60) version of the Hand Laterality Judgement Task. **Panels A and B** show the Reaction Time and Accuracy measures by Rotation Angle and View. **Panel C** shows the ‘biomechanical constraints effect’ (medial vs lateral difference in milliseconds) for the corresponding pairs of angles. **Panel D** shows the ‘biomechanical constraints’ effect collapsing across angles but splitting by View and Laterality.

#### Accuracy

Overall, 22,036 valid trials were analysed. Extreme evidence for including the main effect of Angle (BF_inc_ = 5.95×10^22^), a main effect of View (BF_inc_ = 2.4×10^4^), and their interaction (BF_inc_ = 4.4×10^4^) was found. Extreme evidence against including a main effect of Laterality (BF_01_ = 2.08×10^4^) was found.

The main effect of Angle was driven by accuracy generally decreasing with stimulus rotation (Fig. 4C; see Table 2 for pairwise comparisons). The main effect of View was explained by strong evidence in favour of the palmar views being more accurate than the dorsal views (palmar: 89.78% [85.89, 93.1]; dorsal: 88.4% [84.63, 91.7]; BF_10_ = 18.47). The interaction between Angle and View was explained by the effect of Angle being stronger for dorsal views compared to palmar views at the comparisons 90° vs 135°, 135° vs 180° and 180° vs 225° (dorsal: BF_10_ = 16.72; BF_10_ = 385.21; BF_10_ = 4.94×10^5^; palmar: BF_01_ = 1.49; BF_10_ = 2.05; BF_10_ = 5.03, respectively).

#### Biomechanical constraints effect

The model analysed 14,868 valid trials. There was extreme evidence for including a main effect of Direction (BF_inc_ = 3.34×10^33^), as shown by medial rotations (mean = 1126.22ms [1049.06, 1203.21]) being faster than lateral rotations (mean = 1233.66ms [1155.83, 1310.36]), consistent with the ‘biomechanical constraints’ effect (difference = -107ms [-128, -86.7], BF_10_ = 8.03×10^8^). This effect was critically conditioned on View, as shown by extreme evidence of including a Direction x View interaction (BF_inc_ = 5.09×10^15^), whereby the difference was only present for palmar views (difference = -197.6ms [-226.5, -168.0], BF_10_ = 9.8×10^13^) and not for dorsal views (difference = -17.2ms [-46.7, 11.9], BF_01_ = 11.49). Evidence for including a Direction x Laterality interaction was also extreme (BF_inc_ = 9.61×10^5^), whereby the medial vs lateral difference was larger for right hands (difference = -117ms [-146, - 88.6], BF_10_ = 4.79×10^6^) than for left hands (difference = -98ms [-127, -68.9], BF_10_ = 5.29×10^4^). Extreme evidence for including a three-way interaction (Fig. 4D) was also found (BF_inc_ = 1.23×10^6^), showing that for dorsal views, evidence against biomechanical constraints was stronger for left hands (difference = 2.21ms [-38.9, 43.17), BF_01_ = 17.85) than right hands (difference = -36.74ms [-77.4, 3.99], BF_01_ = 3.76). In the palmar view, the effect was comparable between sides (right = -196.7ms [-238.1, -156.1], BF_10_ = 6.12×10^8^; left = -198.39ms [-239.6, -157.41], BF_10_ = 6.96×10^9^).

#### Effects of Response Mode

The main results are shown in Fig. 5. For reaction time, evidence for including a main effect of Response Mode was extreme (BF_inc_ = 1.66×10^3^). Evidence in favour of equivalence was moderate comparing the Left-hand Group and the Bimanual Group (BF_01_ = 4.51, ROPE = 0 ± 38.9ms). However, the Right-hand Group was slower than the other two groups (Right-hand: 1364.03ms [1234.43, 1491.26]; Left-hand: 1115.60ms [988.68, 1245.60]; Bimanual: 1122.24ms [995.66, 1248.29]), and evidence of non-equivalence was moderate comparing the Right-hand Group and the Bimanual Group (BF_10_ = 9.32), and strong comparing the Right-hand Group and the Left-hand Group (BF_10_ = 11.74). Response Mode only interacted with View (BF_inc_ = 3140), but not with Angle (BF_01_ = 4.51) and Laterality (BF_01_ = 4.51). Extreme evidence against including three-way and four-way interactions was found (BF_01_ > 3.57×10^6^). The interaction between Response Mode and View was driven by evidence of non-equivalence between the groups being stronger (i.e. the Right-hand group responding more slowly) for the dorsal views (Right-hand vs Bimanual: BF_10_ = 28.46; Right-hand vs Left-hand: BF_10_ = 30.42) than the palmar views (Right-hand vs Bimanual: BF_10_ = 2.71; Right-hand vs Left-hand: BF_10_ = 3.69). Evidence for equivalence between the Bimanual and Left-hand group did not change across views (dorsal: BF_01_ = 4.95; palmar: BF_01_ = 4.90).

**Figure 5.**
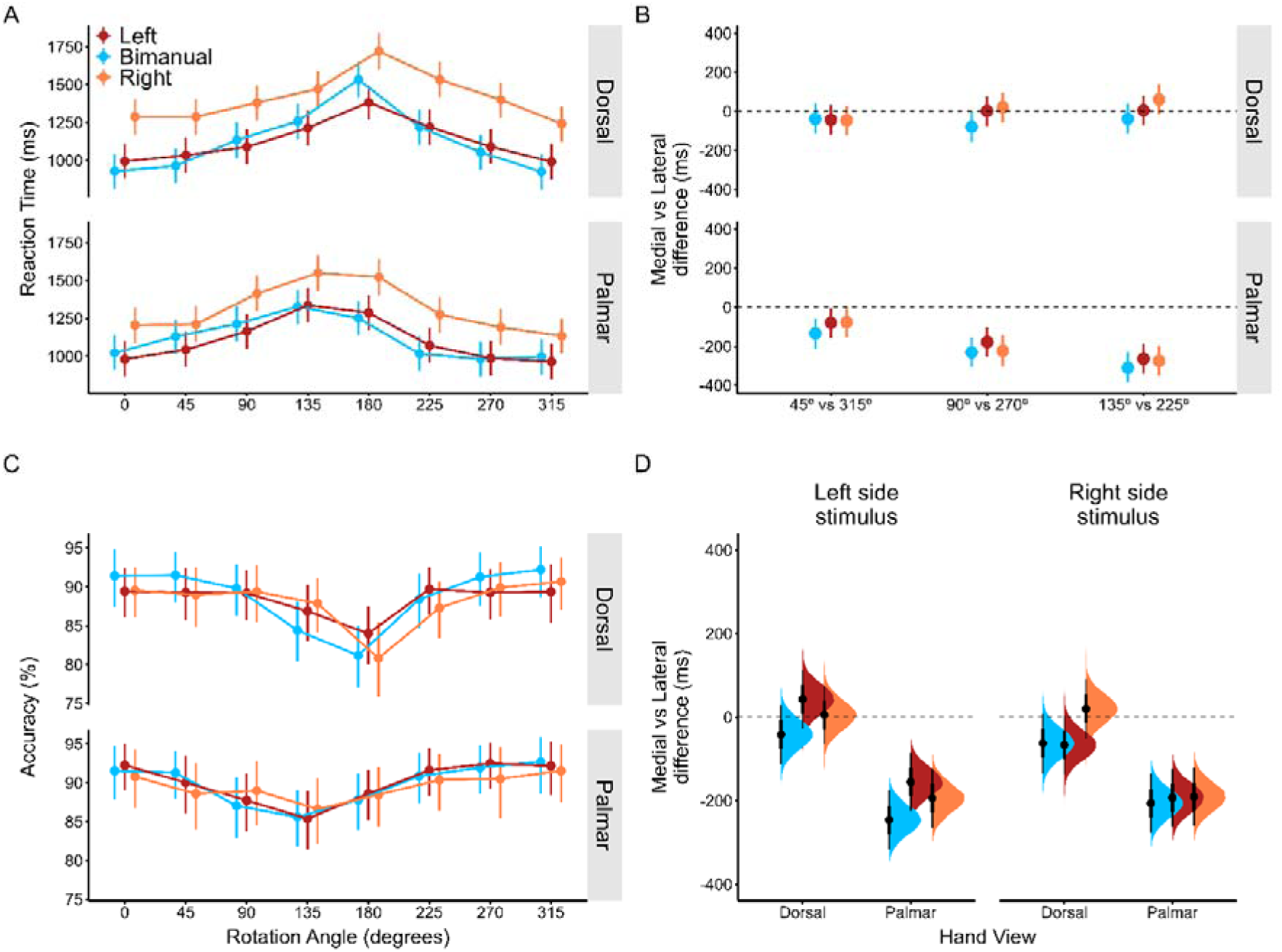
Data from the online version of the Hand Laterality Judgement Task comparing Left-hand, Bimanual and Right-hand Response Modes (N=20 each). **Panels A and B** show the Reaction Time and Accuracy measures by Rotation Angle and View. **Panel C** shows the ‘biomechanical constraints effect’ (medial vs lateral difference in milliseconds) for the corresponding pairs of angles. **Panel D** shows the ‘biomechanical constraints’ effect collapsing across angles but splitting by View and Laterality.

For accuracy, extreme evidence against including a main effect of Response Mode (BF_01_ = 4.08×10^4^) was found. In fact, the overall accuracy was 89.31% [85.56, 92.54] for the Bimanual Group, 89.2% [85.59, 92.34] for the Left-hand group and 88.77% [84.62, 92.32] for the Right-hand Group. Evidence for equivalence was found (Bimanual vs Right-hand: BF_01_ = 15.38; Bimanual vs Left-hand: BF_01_ = 25.64; Right-hand vs Left-hand: BF_01_ = 15.87; ROPE = 0 ± 1.62%). Extreme evidence against including all possible two-way or three-way interactions including Response Mode (BF_01_ > 2.68×10^6^), and the four-way interaction (BF_01_ = 4.59×10^23^) was found.

For the ‘biomechanical constraints’ model, very strong evidence against including a Response Mode x Direction interaction (BF_01_ = 90.91), as well as a three-way interaction with View (BF_01_ = 50) or Laterality (BF_01_ = 1029.87), or a four-way interaction (BF_01_ = 19493.18) was found. The medial vs lateral difference was present in all groups (Left-hand = -92.9ms [-129, -57.5], BF_10_ = 2.9×10^3^; Bimanual = - 139.2ms [-175, -103.9], BF_10_ = 4.69×10^6^; Right-hand = -90.0ms [-126, -54.8], BF_10_ = 1.51×10^3^). The magnitude of the ‘biomechanical constraints effect’ was equivalent between groups, with moderate to strong evidence in favour of the null hypotheses (Bimanual vs Left-hand: BF_01_ = 5.29; Bimanual vs Right-hand: BF_01_ = 4.37; Left-hand vs Right-hand: BF_01_ = 55.56).

### Comparison of in-person and online versions (bimanual groups)

Socio-demographic characteristics were similar across the two Bimanual groups (age: BF_01_ = 2.54; sex: BF_10_ = 1.38), as well as MIQ-3 scores (Total: BF_01_ = 1.17; Kinesthetic: BF_01_ = 2.59; Internal Visual: BF_01_ = 1.73; External Visual: BF_10_ = 2.6). In the HLJT, rejection rates were similar for the online group compared to the in-person group (online = 2.03 ± 3.55%, in-person = 1.54 ± 1.54%, BF_01_ = 2.84). The overall time to complete the task was slightly longer for the online group than the in-person group (online = 21m 30s ± 5m 27.6s, in-person = 17m 42s ± 1m 17.4s; BF_10_ = 9.17).

For reaction time, 13,250 trials were analysed. Very strong evidence against including a main effect of Group (Fig. 6A), and higher-order interactions including this factor was found (all BF_01_ > 5.37×10^4^). Overall, the reaction time was equivalent across the groups (in-person = 1128.35ms [1018.9, 1235.87], online = 1114.05ms [1006.75, 1224.12], BF_01_ = 5.16; ROPE = 0 ± 34.77ms).

**Figure 6.**
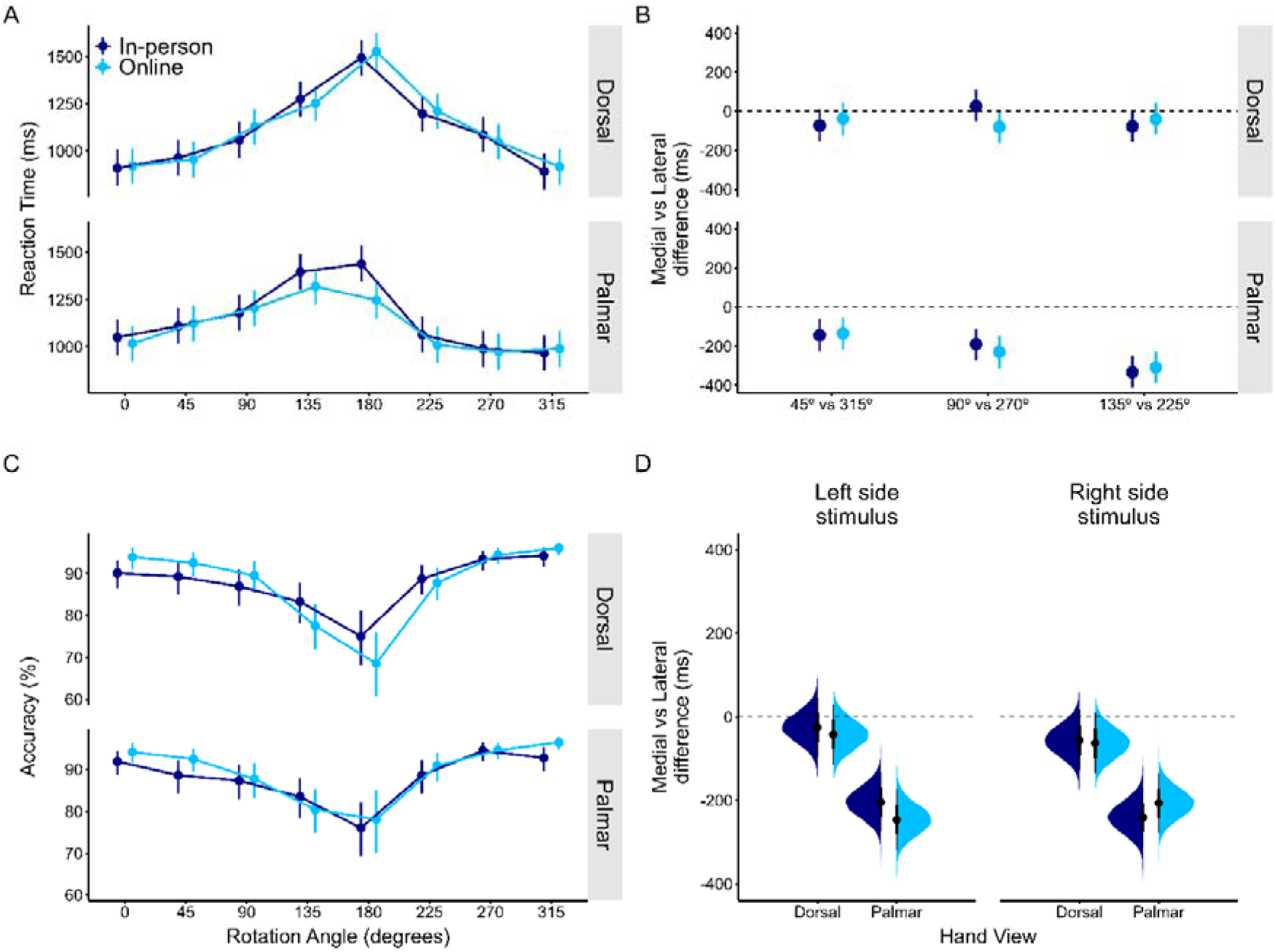
Comparison of the in-person and online Bimanual groups (N=20 each) of the Hand Laterality Judgement Task. **Panels A and B** show the Reaction Time and Accuracy measures by Rotation Angle and View. **Panel C** shows the ‘biomechanical constraints effect’ (medial vs lateral difference in milliseconds) for the corresponding pairs of angles. **Panel D** shows the biomechanical constraints effect collapsing across angles but splitting by View and Laterality.

For accuracy (Fig. 6C), 15,086 trials were analysed. Extreme evidence against including a main effect of Group, and two-way, three-way or four-way interactions including this factor was found (all BF_01_ > 6.06×10^9^). In fact, overall accuracy showed moderate evidence of equivalence (in-person = 87.77% [83.54, 91.37], online = 88.46% [84.5, 91.86], BF_01_ = 6.17; ROPE = 0 ± 1.66%).

For the ‘biomechanical constraints’ effect, 10,142 trials were analysed (Fig. 6D). Extreme evidence against including a Group x Direction interaction was found (BF_01_ = 47.62). In fact, the effect was similar across versions (in-person: -132ms [-166, - 96.9]; online = -139.2ms [-175, -103.9]), with very strong evidence for equivalence (BF_01_ = 34.48; ROPE = 0 ± 32.54). Extreme evidence against including the interactions of Group with View, Laterality, the three-way interaction and the four-way interaction was found (all BF_01_ > 28.57).

## DISCUSSION

This study developed an open-source version of the HLJT for both in-person and online use. The study found that a bimanual Response Mode was largely equivalent to responding with the feet in a lab setting. In addition, the bimanual Response Mode in an online version was practically equivalent to its in-person counterpart. Both versions reproduced the classical behavioural effects of this task, including the ‘biomechanical constraints’ effect. Finally, the comparison between bimanual and unimanual Response Modes showed only a general increase in reaction time in the group responding with the right hand relative to responding bimanually or with the left hand (which were equivalent). This effect was slightly stronger for the dorsal view, did not interact with any other factors of the task, and was not present for accuracy nor affected the ‘biomechanical constraints’ effect.

### Developing a standardized Hand Laterality Judgement Task

Standardisation of key parameters remains an unaddressed challenge in the field of motor imagery, specifically in terms of assessment measures. This not only applies to the HLJT, but a variety of methods (Suica et al., 2022). While we do not advocate for the use of our specific paradigm in all future studies (as some decisions were made based on feasibility or reliability considerations), we hope to partially address this issue. By providing an open-source resource with reasonable default choices but allowing flexibility for future studies (see below), we hope to contribute to creating a common framework for researchers and clinicians interested in using the HLJT.

A wide variety of paradigms have been employed regarding the HLJT. This includes different manipulations of the two main factors of rotation angle and hand view. Some studies have used up to 12 different rotations (i.e. increments of 30°) (Cocksworth & Punt, 2013) or as few as 4 (i.e. increments of 90°) (Conson et al., 2020; Saimpont et al., 2009). Most studies use a number in-between these extremes, usually 6 (de Vries et al., 2013; Ionta et al., 2007) or 8 angles (Brady et al., 2011; Mibu et al., 2020). We decided to use 8 different rotations (i.e. increments of 45°) to maintain the balance between using enough angles to identify a clear slope from 0° to 180°, while reducing the overall number of trials required and therefore the overall time to complete the task. With this paradigm, we could distinctly detect both the main rotation effect for accuracy and reaction time, but also the ‘biomechanical constraints’ effect at each specific rotation, in a task that lasted around 20 minutes overall.

The literature has been highly heterogeneous in the use of different hand views in the HLJT. In the first well-known description of the task, Parsons used only hand drawings of backs and palms of the hands (Parsons, 1987). Subsequently, other studies incremented the variability in this parameter by introducing rotations in the horizontal axis. For example, a ‘thumb’ view or a ‘pinkie’ view have been used, as well as intermediate rotations in-between (Meng et al., 2016; Vannuscorps et al., 2012). In fact, the effect of hand view is still controversial, as most studies suggest that the ‘biomechanical constraints’ effect is only found in the palmar view and not in the dorsal view (Conson et al., 2020; Meng et al., 2016; Mibu et al., 2020), which is clearly consistent with our present results. However, some work suggests that rotation in the horizontal axis is necessary to detect a ‘biomechanical constraints’ effect, which was not observed with simple rotations in the frontal axis in either view (Ter Horst et al., 2010). For the development of our proposed task, we decided to maintain the ‘traditional’ palmar-dorsal paradigm to keep consistency with most of previous studies. This paradigm clearly identified a ‘biomechanical constraints’ effect for the palmar view and found strong evidence for its absence in the dorsal view. We interpret this in line with a growing body of research suggesting that the palm and the back of the hand may be processed using different cognitive strategies, and that only the palm of the hand may trigger motor imagery-based processing (Conson et al., 2021; Nagashima et al., 2019).

Several technical parameters of the HLJT paradigm used in this study were fixed or chosen based on feasibility and reliability. For instance, we decided to include a practice block with all the stimuli to allow the user to familiarise themselves with the task. We consider this will be helpful for future applications, specifically for clinical uses, as it will aid individuals to understand how the task works with minimal explicit supervision by the researcher or clinician, which in our case was necessary for the online version. Regarding the decision of providing feedback on accuracy on a trial-by-trial basis, we chose to include this in the test blocks as we anticipated it would enhance the engagement in the remote version of the task. To maintain consistency across versions, we used the same procedure in the lab. We believe this may be a key parameter to consider in future studies, as it provides participants with online feedback during the task that allows them to adjust their performance if necessary. In addition, we decided to keep the image on-screen until a response was provided, as restricting the time that the participant has to process the stimulus would have limited the generalisability of the paradigm to populations which are generally slower than our sample of young, healthy individuals (e.g. older people or people with neurological disorders). In addition, we decided to use 12 repetitions per unique stimulus. A common rule of thumb in behavioural experiments is to include 8-10 repetitions (Matthews, 2011), but we decided to add more given the remote nature of some of the comparisons, as we expected some participants would not be doing the task as focused as if they were in the lab. Finally, we decided to allow participants to have a break between test blocks and resume the task at their discretion (with a minimum break of 10s). This, in turn, could explain why the online version took longer than the in-person version, as participants may have taken longer breaks in the latter. We made this decision to reduce potential effects of fatigue in future applications and maintain engagement.

We are aware that the specific paradigm used in the present study, though sensible for our purposes, may not meet all the requirements of future uses. For instance, in the clinical field such a task could likely induce fatigue in some individuals and therefore a shorter paradigm might be preferred, whereas a neuroscience study might want to include more rotation angles or different hand views, or might prefer not to provide feedback in the test trials, modify the number of repetitions, etc. Therefore, the task that we publicly share has been modified to allow the user to choose between all these possibilities. At the time of writing, the in-person task has some predefined parameters that can be selected with a mouse click at the beginning of the experiment, including technical considerations such as language (English, Spanish and French are currently supported, and users could add their own translations), whether to include a practice block or not, whether to provide feedback in test blocks or not, the duration of this feedback (0.3s, 0.5s, 0.8s and 1s are available) and the number of repetitions per unique stimulus (12, 8 and 4 are available). Furthermore, the user can select specific parameters of the HLJT, such as the number of rotational angles (4, 6, 8 and 12 are currently available) and the hand views (palmar, dorsal or both). We have set sensible defaults for all these parameters, but changing the defaults is as simple as changing the order of the options in the experiment settings. Furthermore, the range of options for these and other parameters could be easily extended without extensive programming requirements, thanks to PsychoPy2’s capabilities (Peirce et al., 2019). We hope all the above options will allow a wide range of uses in future studies and applications.

### The use of the HLJT in online studies and the use of the hands to respond to the task

Recent advancements in software development have allowed us to leverage the use of online platforms for collecting potentially more representative data, on a larger scale and in a shorter period of time (Bridges et al., 2020; Helms et al., 2021; Johnson et al., 2022). We aimed to translate this development into the field of motor imagery research by testing an online version of the HLJT. Previous work had suggested a confound with responding to the task with the hands would impede to use this Response Mode in the task, as the use of the same effector could interfere with information processing (Cocksworth & Punt, 2013). This would have limited the development of an online HLJT, which requires participants to respond on their own keyboard, and therefore was addressed in this study as part of the development procedure.

Our findings, nonetheless, were consistent with the idea that a manual Response Mode does not meaningfully interfere with this task. In the in-person version, we found evidence for equivalence between feet and bimanual responses in terms of accuracy and the ‘biomechanical constraints’ effect, and evidence against higher-order interactions with all other factors. We only observed moderate evidence for non-equivalence in terms of reaction time (the Bimanual group being approximately 50ms slower than the Foot group across all conditions, without interactions between Response Mode and other factors). We consider this is the main ‘effector confound’ of using the hands to respond, as reaction times are generally faster in manual responses than foot responses in other tasks (Simonen et al., 1995). This effect is consistent with previous work showing longer reaction times for manual responses compared to responding verbally (Cocksworth & Punt, 2013), though they found much larger differences (around 280ms). As they used unimanual Response Modes, which arguably represent a more complex paradigm than responding bimanually, this may explain the different magnitude of the effects. Both studies coincided in the absence of interactions of Response Mode with the rotation angle and hand view, and although Cocksworth & Punt observed lower accuracy for manual responses, the effect was small (difference ≈ 2%). We did not find such an effect (between-group difference < 1%), probably because we used equivalence tests instead of null-hypothesis significance testing. Based on our results, we believe the ‘effector confound’ arguably does not represent a meaningful difference which should prevent to use bimanual responses in the task.

When comparing different bimanual and unimanual Response Modes, we found an unexpected result. The group responding with their right-hand showed generally longer reaction times than the other two groups. This partially contradicts previous findings suggesting that both unimanual Response Modes were similar (Cocksworth & Punt, 2013). Two (potentially interacting) reasons may explain this behaviour. First, responding to this task with two fingers of one hand is arguably more complex than responding with one finger of each hand, which is a more intuitive method. Therefore, a relative slowness in the two unimanual groups was expected compared to the Bimanual group. However, both unimanual groups should have responded similarly, which was not the case. Our (speculative) interpretation relates to the second possible source of this effect; as all participants in our study were right-handed, they most likely were comparing their *dominant* hand to the observed stimuli while making the laterality judgements (i.e. they compared the orientation of the stimulus with their dominant hand, then judged if it could be congruent or not to make the right/left decision). There is previous evidence suggesting this effect may be present in the HLJT (Ní Choisdealbha et al., 2011). In the Left-hand group, this did not generate any conflicts between the processing of the stimulus and the response, as they were responding with their *non-dominant* hand. However, for the Right-hand group, a conflict might have arisen, as the hand used for the visual comparison was the same as the one used to provide the response. The fact that the effect was stronger for the dorsal stimuli further supports this idea, as the dorsal view of the hand is thought to trigger more visual than motor processing, perhaps because we are more familiar with seeing the back of our hands (Bläsing et al., 2013; Zapparoli et al., 2014). Notably, we did not find a corresponding effect for the Bimanual group, who also used their right-hand. We therefore speculate that a possible interaction between these effects may have been present; requiring participants to perform the more complex unimanual version of the task with their right hand may have interfered with the ability to compare their own right hand with the on screen stimulus.

Aside from the findings discussed above, all other comparisons across Response Modes provided strong evidence for equivalence, including all accuracy models and the ‘biomechanical constraints’ effect. While future studies should consider the above-mentioned particularities, we believe that for the purpose of applying the HLJT in applied and online contexts, these findings generally support the use of bimanual and unimanual responses in this task.

### Strengths and Limitations

This study has some limitations. First, we did not investigate the differences between manual and verbal Response Modes, which have been also used in previous studies. We decided to compare a bimanual mode against a foot mode, as these are physically more similar than responding verbally, and this allows to rule out the possibility that slowness due to manual responses can be attributed to the response command simply reaching the effector more quickly. In addition, the feasibility of using hand and foot Response Modes in the task is higher as reaction times from button presses are unambiguous, whereas vocal responses require more training to prevent hesitations or pauses. Similarly, the accuracy of a response from a button press is immediately available allowing rapid feedback, whereas for vocal responses this is not currently possible. Future researchers could build upon our current open-source experiment to implement a vocal response version, if needed.

Second, as part of the study was run fully online, we were not able to assure participants in the unimanual groups were not using the contralateral hand to facilitate performance, as we did not explicitly instruct them what they could or could not do with it. However, given that most evidence indicated equivalence across Response Modes in this version, as well as comparing online and in-person versions, we believe the instructions given were specific enough to obtain reliable estimates in this case. In the future, more explicit guidance could be incorporated.

Third, in the in-person version of the task, the sample was composed of mostly females in the bimanual group, and this can influence HLJT performance (Conson et al., 2020). Therefore, this introduces a potential limitation, as we did not include sex as a covariate in any of the analyses because it was beyond the scope of this paper and we had an unequal distribution between the groups.

Finally, we did not include a control task with non-biological stimuli, to further establish whether the slowdown in processing that was observed is specific to the HLJT or a general pattern also shown in other mental rotation tasks. As we were mostly interested in the ‘biomechanical constraints’ effect, and it would not be present in non-biological stimuli (Bek et al., 2022), we decided not to include such a control task, as it would have doubled the time needed to complete the experiment, potentially compromising the feasibility of the online version of the task.

## CONCLUSIONS

An open-source, freely available Hand Laterality Judgement Task was developed for in-person and online use. The task reproduced established phenomena of this paradigm, both in-person and remotely, and across different Response Modes. For the in-person version, evidence for equivalence between a foot and a bimanual Response Mode was found in terms of accuracy and the ‘biomechanical constraints’ effect. While reaction times were slightly longer in the bimanual group, we found evidence against related higher-order interactions. For the online version, evidence for equivalence between the bimanual and left-hand responses was found for all measures, whereas longer reaction times were found for the Right-hand responses, predominantly for the dorsal view of the hand. Evidence against all other higher-order interactions was found. Evidence of equivalence between the two bimanual groups, in-person and online, was also observed. We conclude that both in-person and online versions reliably replicated key behavioural effects in the HLJT, providing a standardized (but also highly customizable) version of the paradigm that can be readily applied in future studies.

## ACKNOWLEDGEMENTS

We would like to thank Andrés de los Santos Suárez for his assistance with the creation of the figures, and Alfredo Lerín Calvo, Celia López-Cañadillas and Alfredo Hernando Jorge for their help with participant recruitment.

## AUTHOR ROLES (CRediT)

| Author | Conceptualization | Methodology | Software | Validation | Formal Analysis | Investigation | Resources | Data Curation | Writing – original draft | Writing – reviewing & editing | Visualization | Supervision | Project Administration | Funding Acquisition |
| --- | --- | --- | --- | --- | --- | --- | --- | --- | --- | --- | --- | --- | --- | --- |
| MMV | X | X | X | X | X | X |  | X | X | X | X |  | X |  |
| SMcA |  | X | X |  | X |  |  |  |  | X | X |  |  |  |
| BMW |  |  | X |  | X |  |  |  |  | X |  |  |  |  |
| EVC |  |  |  |  |  |  |  |  |  | X |  |  |  |  |
| RMH | X | X |  | X |  |  | X |  | X | X |  | X | X | X |

## SUPPLEMENTARY MATERIALS

### Details of Bayesian models

For accuracy, the zero-and-one inflated beta-regression models were modelled with by-group (i.e. Response Mode) and by-Angle random intercepts and slopes for the parameters ‘phi’, ‘zoi’ and ‘coi’, which estimated the measures of variability in the beta distribution part of the model, the proportion of 0s and 1s in the distribution, and the proportion of 1s within that proportion, respectively. This allowed the models to converge better. These parameters were not used for any inferences, as the most relevant measure was the beta distribution representing the continuous probability of a correct response (i.e. Accuracy), which was modelled as explained in the Methods section of this paper.

### Customisability options in the Hand Laterality Judgement Task

Both local and online versions of the HLJT were developed in a modular manner, that meaning that the level of flexibility is relatively high. The details about customisation are fully described in a ‘Readme’ file specific to each version. As a general description, in the current versions the experimenter can select the following parameters:

- Language: English (default), Spanish and French. Language to display instructions and messages throughout the task.
- Response mode: Both hands (default), Right hand or Left hand. Whether the participant should respond bimanually or unimanually.
- Practice block: Yes (default) or No. Whether a practice block with all the stimuli should be displayed before the actual test blocks.
- Number of angles: 4 angles with increments of 90° (default), 6 angles with increments of 60°, 8 angles with increments of 45° or 12 angles with increments of 30°.
- Hand views: Palmar and Dorsal (default), Palmar or Dorsal. Whether the stimuli should appear in one hand view only or both.
- Number of repetitions: 12 (default), 8 or 4. Number of repetitions per unique stimulus in the test blocks overall (i.e. in total excluding the practice block). As of now the number of test blocks is fixed at 4, therefore, the total number of repetitions must be a multiple of 4.
- Feedback: 0.3 seconds (default), 0.5 seconds, 0.8 seconds, 1 second or No feedback. Time of feedback for accuracy (correct/incorrect) on a trial-by-trial basis. This only applies to test blocks. If a practice block is included in the experiment, feedback throughout this block is always provided.

In these versions, the inter-trial interval (ITI) for both the practice and test blocks, is set at a random number between 0.6 and 1 seconds, with more probability in the range 0.75 to 0.85 seconds, with the goal of having a random ITI throughout the experiment with an average duration of ∼0.8 seconds every 50 trials.

**Supplementary Tables with descriptives statistics in the in-person version.**

**Supplementary Table 1.** Mean and SD for response time (milliseconds) for each Response Mode (group) according to Angle (0° to 315°), Laterality (left or right) and View (dorsal or palmar).

| Angle | Group | Left |  |  |  | Right |  |  |  |
| --- | --- | --- | --- | --- | --- | --- | --- | --- | --- |
|  |  | Dorsal |  | Palmar |  | Dorsal |  | Palmar |  |
|  |  | Mean | SD | Mean | SD | Mean | SD | Mean | SD |
| 0° | Foot | 827.92 | 253.69 | 1017.97 | 423.06 | 793.29 | 278.66 | 904.85 | 347.17 |
|  | Bimanual | 978.47 | 343.97 | 1126.10 | 481.00 | 924.41 | 414.39 | 1070.51 | 467.73 |
| 45° | Foot | 910.84 | 349.62 | 1022.03 | 404.03 | 855.49 | 303.41 | 1001.50 | 385.67 |
|  | Bimanual | 1081.33 | 470.16 | 1179.46 | 450.01 | 970.79 | 436.83 | 1152.25 | 444.00 |
| 90° | Foot | 1079.28 | 407.96 | 1187.59 | 465.03 | 1005.50 | 387.16 | 1184.31 | 438.60 |
|  | Bimanual | 1256.92 | 498.70 | 1410.35 | 532.55 | 1261.10 | 543.17 | 1369.96 | 552.18 |
| 135° | Foot | 1328.12 | 538.27 | 1195.54 | 497.79 | 1318.98 | 505.93 | 1046.16 | 423.22 |
|  | Bimanual | 1531.75 | 573.52 | 1492.67 | 617.45 | 1504.95 | 598.91 | 1311.45 | 523.80 |
| 180° | Foot | 1084.28 | 415.45 | 977.93 | 409.21 | 1033.37 | 495.20 | 864.78 | 359.97 |
|  | Bimanual | 1230.12 | 453.12 | 1116.48 | 433.89 | 1171.71 | 460.69 | 1010.36 | 380.81 |
| 225° | Foot | 964.34 | 374.04 | 869.37 | 312.95 | 866.95 | 373.05 | 822.56 | 334.15 |
|  | Bimanual | 1119.10 | 419.03 | 1000.62 | 370.66 | 1029.21 | 426.37 | 963.80 | 365.45 |
| 270° | Foot | 829.88 | 286.18 | 871.26 | 322.64 | 773.60 | 301.17 | 799.48 | 320.38 |
|  | Bimanual | 907.83 | 315.06 | 993.13 | 369.08 | 848.45 | 369.50 | 926.59 | 325.69 |
| 315° | Foot | 827.92 | 253.69 | 1017.97 | 423.06 | 793.29 | 278.66 | 904.85 | 347.17 |
|  | Bimanual | 978.47 | 343.97 | 1126.10 | 481.00 | 924.41 | 414.39 | 1070.51 | 467.73 |

**Supplementary Table 2.** Mean and SD for accuracy (%) for each Response Mode (group) according to Angle (0° to 315°), Laterality (left or right) and View (dorsal or palmar).

| Angle | Group | Left |  |  |  | Right |  |  |  |
| --- | --- | --- | --- | --- | --- | --- | --- | --- | --- |
|  |  | Dorsal |  | Palmar |  | Dorsal |  | Palmar |  |
|  |  | Mean | SD | Mean | SD | Mean | SD | Mean | SD |
| 0° | Foot | 89.54 | 30.67 | 86.50 | 34.25 | 93.33 | 25.00 | 88.75 | 31.66 |
|  | Bimanual | 86.25 | 34.51 | 91.67 | 27.70 | 88.75 | 31.66 | 90.76 | 29.03 |
| 45° | Foot | 92.44 | 26.50 | 89.54 | 30.67 | 91.25 | 28.32 | 87.39 | 33.26 |
|  | Bimanual | 90.25 | 29.72 | 87.87 | 32.72 | 87.45 | 33.20 | 86.55 | 34.19 |
| 90° | Foot | 86.44 | 34.31 | 88.14 | 32.41 | 88.19 | 32.35 | 84.68 | 36.09 |
|  | Bimanual | 89.41 | 30.84 | 90.72 | 29.08 | 83.97 | 36.77 | 81.20 | 39.16 |
| 135° | Foot | 77.54 | 41.82 | 79.65 | 40.34 | 71.13 | 45.41 | 78.06 | 41.47 |
|  | Bimanual | 90.68 | 29.14 | 85.71 | 35.07 | 75.95 | 42.83 | 80.95 | 39.35 |
| 180° | Foot | 72.53 | 44.73 | 75.00 | 43.39 | 57.38 | 49.56 | 79.91 | 40.15 |
|  | Bimanual | 76.96 | 42.20 | 74.01 | 43.96 | 69.91 | 45.97 | 75.11 | 43.33 |
| 225° | Foot | 82.35 | 38.20 | 87.39 | 33.26 | 83.97 | 36.77 | 93.64 | 24.45 |
|  | Bimanual | 88.84 | 31.55 | 90.21 | 29.78 | 84.52 | 36.25 | 89.54 | 30.67 |
| 270° | Foot | 91.21 | 28.37 | 93.16 | 25.29 | 89.12 | 31.20 | 95.82 | 20.06 |
|  | Bimanual | 93.31 | 25.05 | 94.58 | 22.68 | 90.00 | 30.06 | 95.00 | 21.84 |
| 315° | Foot | 93.70 | 24.35 | 94.56 | 22.73 | 93.70 | 24.35 | 96.22 | 19.12 |
|  | Bimanual | 92.47 | 26.45 | 92.08 | 27.06 | 93.72 | 24.30 | 92.44 | 26.50 |

**Supplementary Tables with descriptives statistics in the online version.**

**Supplementary Table 3.** Mean and SD for response time (milliseconds) for each Response Mode (group) according to Angle (0° to 315°), Laterality (left or right) and View (dorsal or palmar).

| Angle | Group | Left |  |  |  | Right |  |  |  |
| --- | --- | --- | --- | --- | --- | --- | --- | --- | --- |
|  |  | Dorsal |  | Palmar |  | Dorsal |  | Palmar |  |
|  |  | Mean | SD | Mean | SD | Mean | SD | Mean | SD |
| 0° | Bimanual | 962.59 | 406.00 | 1032.30 | 414.57 | 858.16 | 337.01 | 969.95 | 395.07 |
|  | Left-hand | 1028.31 | 387.03 | 1003.42 | 412.56 | 941.34 | 463.53 | 932.16 | 360.72 |
|  | Right-hand | 1287.04 | 511.94 | 1188.90 | 500.35 | 1222.52 | 522.32 | 1225.30 | 499.89 |
| 45° | Bimanual | 965.95 | 440.92 | 1133.96 | 457.16 | 902.99 | 394.37 | 1085.00 | 518.76 |
|  | Left-hand | 1086.17 | 499.99 | 1100.83 | 431.18 | 955.33 | 426.05 | 962.00 | 386.29 |
|  | Right-hand | 1293.51 | 552.02 | 1201.71 | 495.81 | 1245.36 | 533.03 | 1189.58 | 517.99 |
| 90° | Bimanual | 1118.28 | 574.84 | 1180.72 | 469.77 | 1030.31 | 478.95 | 1152.42 | 550.65 |
|  | Left-hand | 1133.55 | 462.22 | 1187.68 | 501.55 | 1030.35 | 434.90 | 1117.57 | 494.69 |
|  | Right-hand | 1397.70 | 593.83 | 1361.04 | 588.06 | 1315.25 | 562.52 | 1387.97 | 620.70 |
| 135° | Bimanual | 1219.58 | 500.47 | 1312.56 | 539.48 | 1162.24 | 496.69 | 1235.60 | 510.30 |
|  | Left-hand | 1193.48 | 502.08 | 1355.50 | 539.56 | 1189.21 | 500.41 | 1297.34 | 565.08 |
|  | Right-hand | 1441.74 | 551.71 | 1470.49 | 607.11 | 1452.80 | 574.97 | 1458.47 | 602.75 |
| 180° | Bimanual | 1513.65 | 521.88 | 1249.11 | 531.03 | 1496.75 | 487.37 | 1195.21 | 531.28 |
|  | Left-hand | 1330.55 | 505.66 | 1304.30 | 473.24 | 1412.99 | 518.96 | 1251.08 | 493.75 |
|  | Right-hand | 1740.33 | 561.68 | 1547.38 | 626.33 | 1668.54 | 507.88 | 1406.76 | 625.81 |
| 225° | Bimanual | 1272.35 | 506.11 | 1007.86 | 341.32 | 1120.06 | 452.76 | 993.53 | 421.90 |
|  | Left-hand | 1282.56 | 552.32 | 1122.52 | 464.69 | 1137.60 | 466.82 | 1007.20 | 415.26 |
|  | Right-hand | 1521.30 | 543.82 | 1251.39 | 585.67 | 1503.63 | 545.24 | 1194.16 | 522.04 |
| 270° | Bimanual | 1073.38 | 402.84 | 971.10 | 386.32 | 1015.00 | 417.41 | 958.45 | 424.24 |
|  | Left-hand | 1177.24 | 472.97 | 1051.99 | 406.00 | 990.64 | 414.94 | 905.69 | 339.61 |
|  | Right-hand | 1394.91 | 518.85 | 1188.44 | 524.60 | 1409.68 | 544.31 | 1173.47 | 484.47 |
| 315° | Bimanual | 954.32 | 401.12 | 991.58 | 380.06 | 872.23 | 382.46 | 973.24 | 433.82 |
|  | Left-hand | 1082.04 | 507.01 | 970.98 | 384.74 | 874.04 | 352.48 | 926.03 | 384.57 |
|  | Right-hand | 1235.62 | 531.45 | 1088.52 | 477.32 | 1245.80 | 520.03 | 1164.92 | 501.86 |

**Supplementary Table 4.**
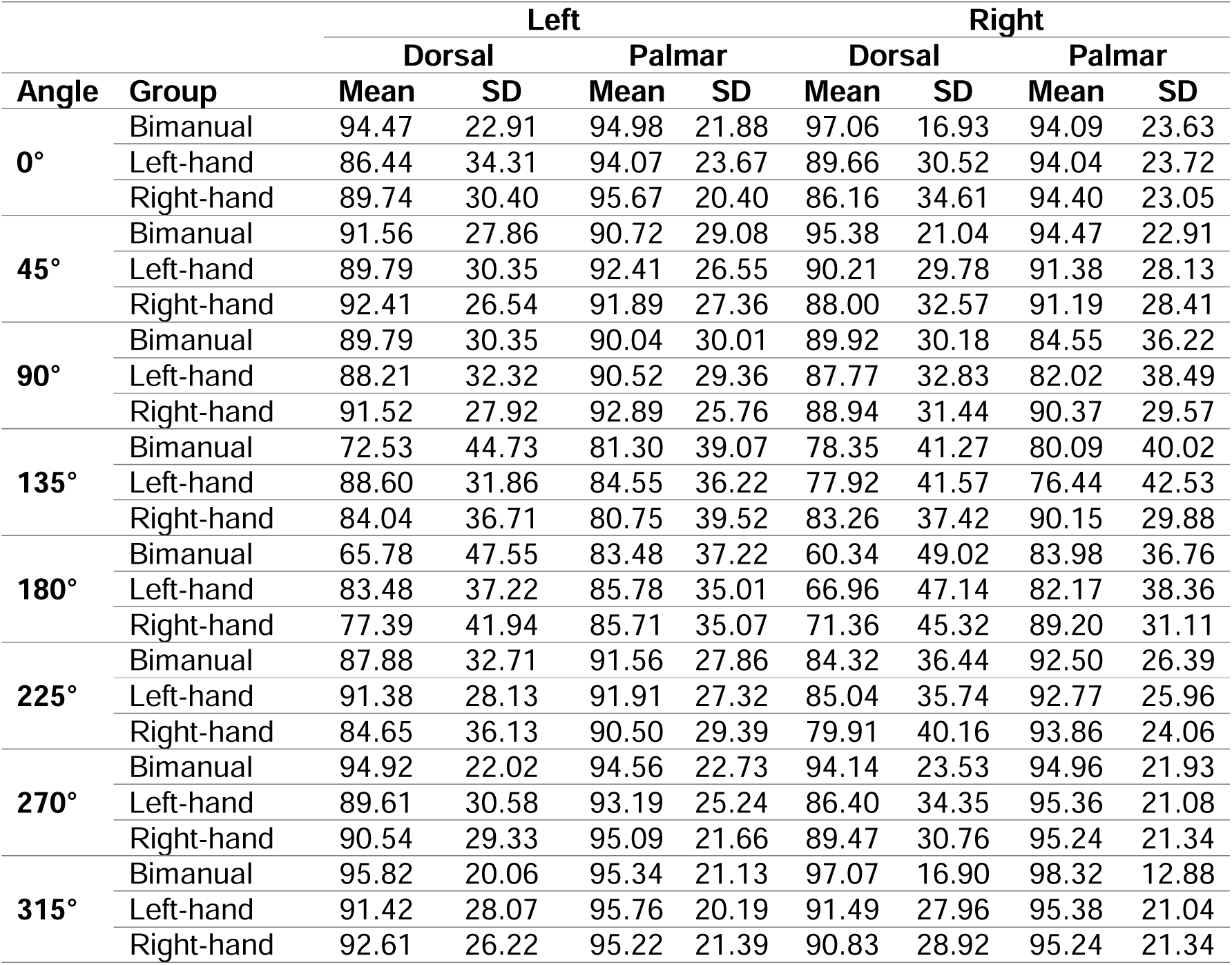
Mean and SD for accuracy (%) for each Response Mode (group) according to Angle (0° to 315°), Laterality (left or right) and View (dorsal or palmar).

| Angle | Group | Left |  |  |  | Right |  |  |  |
| --- | --- | --- | --- | --- | --- | --- | --- | --- | --- |
|  |  | Dorsal |  | Palmar |  | Dorsal |  | Palmar |  |
|  |  | Mean | SD | Mean | SD | Mean | SD | Mean | SD |
| 0° | Bimanual | 94.47 | 22.91 | 94.98 | 21.88 | 97.06 | 16.93 | 94.09 | 23.63 |
|  | Left-hand | 86.44 | 34.31 | 94.07 | 23.67 | 89.66 | 30.52 | 94.04 | 23.72 |
|  | Right-hand | 89.74 | 30.40 | 95.67 | 20.40 | 86.16 | 34.61 | 94.40 | 23.05 |
| 45° | Bimanual | 91.56 | 27.86 | 90.72 | 29.08 | 95.38 | 21.04 | 94.47 | 22.91 |
|  | Left-hand | 89.79 | 30.35 | 92.41 | 26.55 | 90.21 | 29.78 | 91.38 | 28.13 |
|  | Right-hand | 92.41 | 26.54 | 91.89 | 27.36 | 88.00 | 32.57 | 91.19 | 28.41 |
| 90° | Bimanual | 89.79 | 30.35 | 90.04 | 30.01 | 89.92 | 30.18 | 84.55 | 36.22 |
|  | Left-hand | 88.21 | 32.32 | 90.52 | 29.36 | 87.77 | 32.83 | 82.02 | 38.49 |
|  | Right-hand | 91.52 | 27.92 | 92.89 | 25.76 | 88.94 | 31.44 | 90.37 | 29.57 |
| 135° | Bimanual | 72.53 | 44.73 | 81.30 | 39.07 | 78.35 | 41.27 | 80.09 | 40.02 |
|  | Left-hand | 88.60 | 31.86 | 84.55 | 36.22 | 77.92 | 41.57 | 76.44 | 42.53 |
|  | Right-hand | 84.04 | 36.71 | 80.75 | 39.52 | 83.26 | 37.42 | 90.15 | 29.88 |
| 180° | Bimanual | 65.78 | 47.55 | 83.48 | 37.22 | 60.34 | 49.02 | 83.98 | 36.76 |
|  | Left-hand | 83.48 | 37.22 | 85.78 | 35.01 | 66.96 | 47.14 | 82.17 | 38.36 |
|  | Right-hand | 77.39 | 41.94 | 85.71 | 35.07 | 71.36 | 45.32 | 89.20 | 31.11 |
| 225° | Bimanual | 87.88 | 32.71 | 91.56 | 27.86 | 84.32 | 36.44 | 92.50 | 26.39 |
|  | Left-hand | 91.38 | 28.13 | 91.91 | 27.32 | 85.04 | 35.74 | 92.77 | 25.96 |
|  | Right-hand | 84.65 | 36.13 | 90.50 | 29.39 | 79.91 | 40.16 | 93.86 | 24.06 |
| 270° | Bimanual | 94.92 | 22.02 | 94.56 | 22.73 | 94.14 | 23.53 | 94.96 | 21.93 |
|  | Left-hand | 89.61 | 30.58 | 93.19 | 25.24 | 86.40 | 34.35 | 95.36 | 21.08 |
|  | Right-hand | 90.54 | 29.33 | 95.09 | 21.66 | 89.47 | 30.76 | 95.24 | 21.34 |
| 315° | Bimanual | 95.82 | 20.06 | 95.34 | 21.13 | 97.07 | 16.90 | 98.32 | 12.88 |
|  | Left-hand | 91.42 | 28.07 | 95.76 | 20.19 | 91.49 | 27.96 | 95.38 | 21.04 |
|  | Right-hand | 92.61 | 26.22 | 95.22 | 21.39 | 90.83 | 28.92 | 95.24 | 21.34 |

